# Identification and characterization of CD4+ T cell epitopes after Shingrix vaccination

**DOI:** 10.1101/2020.07.29.227082

**Authors:** Hannah Voic, Rory D. de Vries, John Sidney, Paul Rubiro, Erin Moore, Elizabeth Phillips, Simon Mallal, Brittany Schwan, Daniela Weiskopf, Alessandro Sette, Alba Grifoni

**Author notes:** Alessandro Sette and Alba Grifoni contributed equally to this work. Address correspondence to Alba Grifoni.

## Abstract

Infections with varicella zoster virus (VZV), a member of the Herpesviridae family, are associated with a range of clinical manifestations. Primary infection with VZV causes chicken pox, and due to the virus’s capacity to remain latent in neurons, it can reactivate later in life causing herpes zoster (HZ), also known as shingles. Two different vaccines have been developed to prevent HZ, one based on a live attenuated VZV strain (Zostavax) and the other on adjuvanted gE recombinant protein (Shingrix). While Zostavax efficacy wanes with age, Shingrix protection retains its efficacy in elderly subjects (80 years of age and beyond). In this context, it is of much interest to understand if there is a role for T cell immunity in differential clinical outcome, and if there is a correlate of protection between T cell immunity and Shingrix efficacy. In this study, we characterized Shingrix specific ex vivo CD4 T cell responses in the context of natural exposure and HZ vaccination using pools of predicted epitopes. We show that T cell reactivity following natural infection and Zostavax vaccination dominantly targets non-structural proteins (NS), while Shingrix vaccination redirects dominant reactivity to target gE. We mapped the gE-specific responses following Shingrix vaccination to 89 different gE epitopes, 34 of which accounted for 80% of the response. Using antigen presentation assays and single HLA molecule transfected lines, we experimentally determined HLA restrictions for 94 different donor/peptide combinations. Finally, we used our results as a training set to assess strategies to predict restrictions based on measured or predicted HLA binding and the corresponding HLA types of responding subjects.

**Importance:** Understanding the T cell profile associated with the protection observed in elderly vaccinees following Shingrix vaccination is relevant to the general definition of correlates of vaccine efficacy. Our study enables these future studies by clarifying patterns of immunodominance associated with Shingrix vaccination, as opposed to natural infection or Zostavax vaccination. Identification of epitopes recognized by Shingrix-induced CD4 T cells and their associated HLA restrictions enables the generation of tetrameric staining reagents and, more broadly, the capability to characterize specificity, magnitude and phenotype of VZV specific T cells.

## Introduction

VZV, or HHV3, is a dsDNA virus of about 125kb and belongs to the Herpesviridae family. The corresponding proteome encompasses 69 proteins (13), of which 37 are structural, including Envelope, Tegument, and Capsid, and 32 are non-structural proteins involved in multiple functions, such as viral replication, innate host immunity evasion, and transcriptional regulation connected to the viral latency state (13). Three protein layers comprise a VZV particle: a nucleocapsid containing dsDNA genome, a tegument layer, and an envelope lipid bilayer containing viral glycoproteins (12).

Primary infection generally occurs in children prior to adolescence in temperate climates, but can occur at any age range, and clinically manifests as Varicella or chickenpox, a mild-to moderate disease with symptoms appearing 10-21 days after exposure (4). The characteristic symptom of varicella is the pruritic vesicular rash appearing on the head, torso, and extremities accompanied by additional symptoms including headache, loss of appetite, tiredness and fever (4). The host immune system controls, but does not eliminate, the virus, as VZV enters a life-long latency state mainly located in the sensory neurons of the trigeminal and dorsal root ganglia, but also in other sensory neurons, as well as autonomic, and other cranial nerve and enteric ganglia (12). The reactivation of latent virus is the cause of a second disease state of Herpes Zoster (HZ or Shingles), characterized by a more localized infection (12, 41, 64). Post-herpetic neuralgia (PHN) is a very common neurological complication of HZ that can persist for years, and occurs in ∼15% of cases. Age is the most important risk factor for PHN, with the risk increasing rapidly after 50 years of age (19). It has been estimated that half of the subjects reaching age 85 will experience Herpes Zoster (HZ) reactivation at least once in their lifetime (4). While this estimation may change depending on the population studied, overall this highlights the need for efficient control of this disease.

The mechanism of protection from HZ reactivation is unclear, but cellular mediated immunity (CMI) has been shown to be integral to HZ prevention. Additionally, HZ severity has been clinically observed to be associated with a reduced magnitude of VZV specific effector and effector memory T cells (33).

Three vaccines are currently licensed in the United States for use in preventing VZV associated diseases. Varivax, a live attenuated vaccine for varicella immunity, was first licensed in the United States in 1995, and is part of the recommended childhood vaccine schedule. In 2005, two doses of this vaccine during childhood were recommended in the USA (4). Two vaccines have been developed to protect against Herpes Zoster (HZ): Zostavax, a live attenuated vaccine derived from the Oka virus strain, and Shingrix, a vaccine based on recombinant gE protein vaccination. Zostavax was licensed in 2006 for people 60 years and older (4). Several studies have evaluated the efficacy of Zostavax in preventing HZ, with some initially hypothesizing that the response was reduced as a function of the age of the vaccinee. Indeed, an age effect ranging from 70% efficacy in ages 50 – 59 to a mere 18% efficacy in vaccinees 80 years old or older, has been noted. Further, more recent studies show an overall waning of immunity after 4 years, regardless of age (17), and a very recent study performed in Sweden showed that the effectiveness of Zostavax was reduced to 34% and protection, observed in the 61-75 years age group, was largely absent in individuals >75 years older (2). Overall, Zostavax vaccination was shown to be significantly less effective than Shingrix vaccination, and for this reason, starting July 2020, Zostavax is no longer available (33, 35). The Shingrix vaccine was recently licensed for adults 50 years and older and efficacy trials in adults have demonstrated protection regardless of age, with 91% efficacy observed in vaccinees 80 years or older (27). The Shingrix efficacy at older ages is yet to be fully understood, and additional studies dissecting the responses triggered by this vaccination are important for developing successful vaccination against other pathogens in the elderly population.

The recombinant VZV gE protein is co-formulated with the novel ASB01 adjuvant to induce both a strong CD4+ T cell and gE-specific antibody response (6, 27). VZV-specific CD4+ T cell responses are associated with positive vaccine outcomes (20, 54, 59, 62). The reasons for the remarkable difference in Shingrix efficacy may involve a capacity for eliciting a stronger and/or more focused response against the gE antigen, a broader epitope repertoire or an intrinsic difference in the quality and durability of the responses elicited. Several studies have indicated changes in the quality of the specific CD4+ T cell response towards a predominantly and persistently polyfunctional response after vaccination (7, 27, 34).

While studies have been conducted to assess the immunogenicity of different VZV proteins (32), and the immunological responses to VZV vaccinations have been described elsewhere (31, 36, 59, 60), there is still little known regarding the epitopes recognized by T cells after VZV vaccination. Identifying immunogenic epitopes and designing tetramers specific to VZV epitopes will allow further characterization of the observed polyfunctional response in Shingrix vaccinees and may help establish correlates of protection for HZ patients.

Here we assess the repertoire of CD4+ T cell responses induced by Shingrix vaccination and define a set of epitopes broadly recognized in these vaccinees. Further, we use both predictive binding methods and reductive HLA restriction assays with identified VZV-specific CD4+ T cell epitopes to identify potential tetrameric reagents and immuno-monitoring strategies to address mechanisms of cell-mediated immunity and effective vaccination strategies.

## Results

### Recruitment of childhood exposure, post-shingles, and Zostavax and Shingrix vaccinees donor cohorts

To characterize CD4+ T cell responses following Shingrix vaccination we enrolled 18 Shingrix vaccinees. These vaccinees were further sub-classified depending on their status before Shingrix vaccination as: Shingrix only (n=10), Shingrix after shingles (n=2), and Shingrix after Zostavax (n=5); an additional donor recently vaccinated with Shingrix, but who was previously vaccinated with Zostavax and had developed shingles, was also included. To put our observations in a broader context of other HZ related conditions, we also enrolled three independent cohorts of donors to represent individuals with childhood exposure (n=18), Zostavax vaccinees (n=18), and clinically diagnosed shingles with lack of vaccination (in childhood or otherwise) (n=18). The childhood exposure cohort encompassed donors who, based on date of birth, did not receive childhood vaccination and were likely to have experienced clinical chickenpox; as such they represent the natural exposure to VZV and never experienced shingles. A total of 72 donors participated in this study. Ethnicity, gender, average age, and time from most recent clinical event (Shingrix or Zostavax vaccination or HZ reactivation) are summarized in **Table 1**. The cohorts enrolled herein are based on a pool of volunteers in the San Diego area. Therefore, despite the efforts to enroll a cohort as balanced as possible in terms of gender, ethnicity and age-range we still observed some biases. A significant difference in age is observed across cohorts (p<0.0001 Kruskal-Wallis test), but no significant difference in sex. However, a generalized bias towards male gender and Caucasian’s is observed overall. Because Zostavax vaccination is not available as of July 2020, and recommendation towards Shingrix vaccination has been in place since early 2019, it was not possible to analyze the two HZ vaccinations at the same time. This has, as a result, generated a significant difference between time of exposure and sample collection between Zostavax and Shingrix vaccinations. Nonetheless, the Zostavax and Shingles cohorts were analyzed after a similar time frame.

**Table 1.**
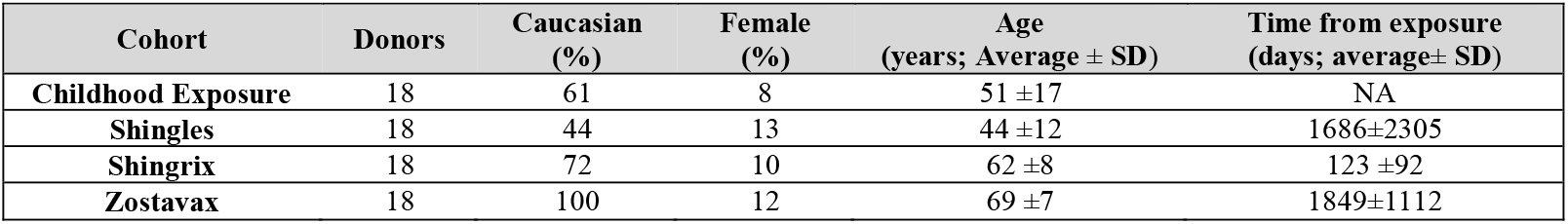
General characteristics of donor cohorts.

| Cohort | Donors | Caucasian (%) | Female (%) | Age (years; Average ± SD) | Time from exposure (days; average± SD) |
| --- | --- | --- | --- | --- | --- |
| Childhood Exposure | 18 | 61 | 8 | 51 ±17 | NA |
| Shingles | 18 | 44 | 13 | 44 ±12 | 1686±2305 |
| Shingrix | 18 | 72 | 10 | 62 ±8 | 123 ±92 |
| Zostavax | 18 | 100 | 12 | 69 ±7 | 1849±1112 |

### General features of VZV responses in the various cohorts

The study of responses in different settings and geographical locations is important in the context of vaccination targeting different human populations, as different HLA alleles, influencing T-cell immune responses, might be prevalent in different populations. Additionally, to be able to analyze unaltered T cell phenotypes, it is important to study T cell reactivity ex-vivo. However, testing T cell responses ex vivo against each single peptide, while ensuring comprehensive protein and HLA coverage, requires a volume of blood not often available in for human studies. To meet these challenges, we have previously developed the megapool approach, based on pooling large numbers of peptides and formulated to consider sequential lyophilization. These “megapools” (MP) have been used in multiple studies in the context of a number of different indications, from allergy to infectious diseases and vaccination (8, 23, 25, 28, 39).

Shingrix vaccination is based on recombinant gE protein only, while in the context of natural exposure, as well as after a live-attenuated vaccination, an individual is exposed to the entire VZV proteome. To address the differences in terms of T cell reactivity across the different cohorts, we designed megapools targeting the gE protein specifically versus the rest of the VZV proteome. In the case of the gE protein, 123 peptides (15-mers overlapping by 10 residues) spanning the gE protein were arranged into a single “megapool”. The gE protein sequence of the Dumas strain is 100% identical with the gE protein of the Oka strain used in Zostavax.

An additional 302 peptides from the remaining non-gE proteins, derived from the entire VZV proteome, were investigated to identify CD4+ T cell reactivity outside the gE protein. These included 44 epitopes from the published literature curated in the Immune Epitope Database (IEDB, www.IEDB.org)(63), and 258 predicted HLA class II dominant epitopes from the Dumas strain reference sequence (52). To ensure that the pools of peptides tested were based on a similar number of peptides, the non gE peptides were arranged into two different megapools, with one corresponding roughly to structural proteins (S, but excluding gE), and the other to the non-structural proteins (NS), as previously described (18). The full list of peptides contained in the non-gE Megapools (S and NS), as well as additional information regarding the corresponding peptide start position, protein of provenance, and ORF of origin is provided in **Table S1**.

VZV CD4+T cell responses in natural infection and following HZ vaccination have been shown to elicit a broad spectrum of cytokines (5, 9, 36). This suggests that a more comprehensive, cytokine-independent, strategy is necessary to dissect CD4+T cell specific responses across cohorts is required to be able to compare the antigens recognized. For this purpose, we applied a T cell receptor dependent Activation Induced Marker (AIM) assay (10, 26) by analyzing the combined membrane markers for expression of CD137 (41BB) and OX40 molecules within the T cell’s CD4+ compartment after 24 hours stimulation with the gE, S and NS megapools (MP). **Fig 1** shows in detail the gating strategy approach applied.

**FIG 1.**
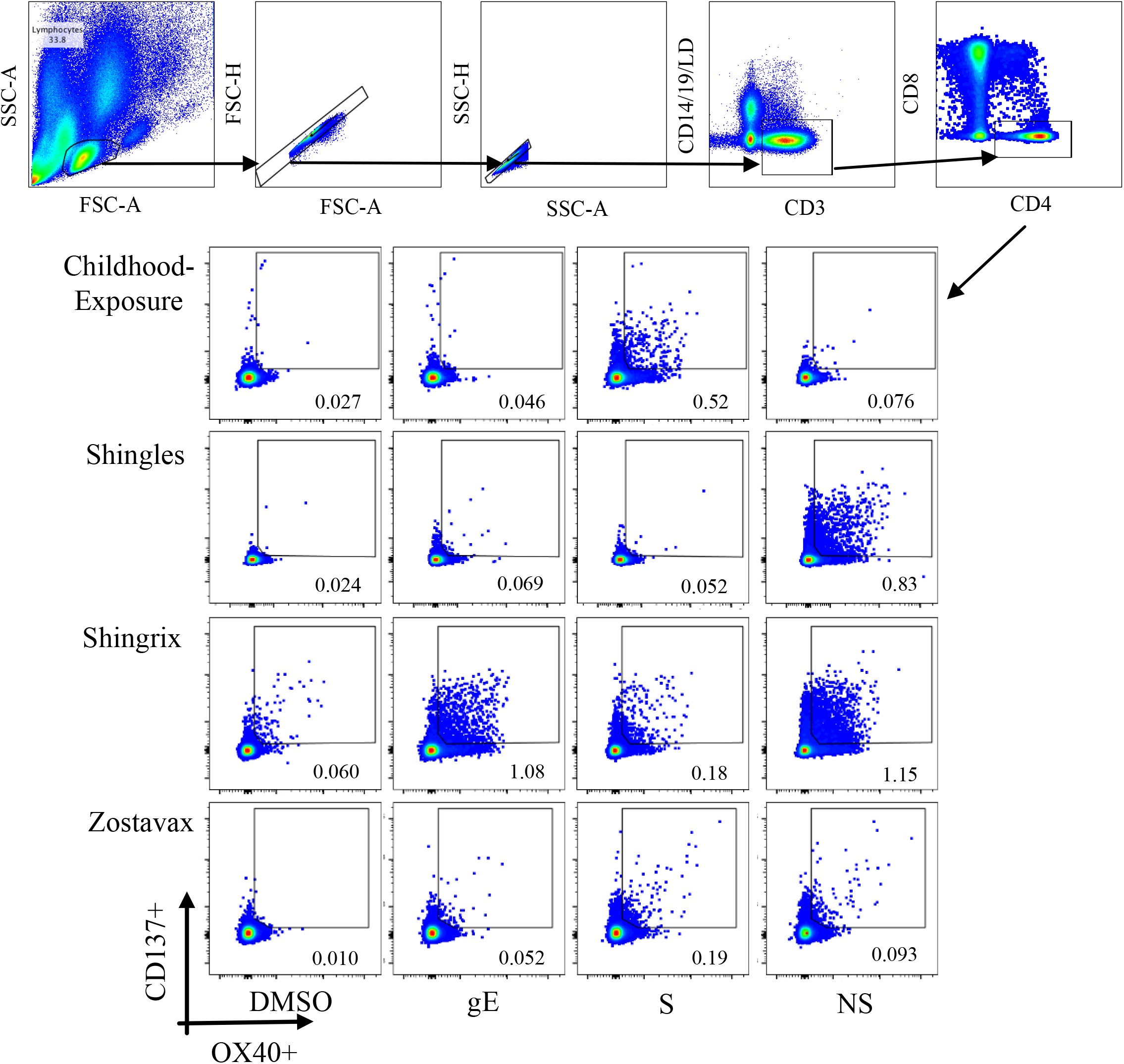
Representative Gating Strategy.

Strong responses to NS proteins were noted in the case of the childhood exposure and Shingles cohorts (p=0.0004 and p<0.0001, respectively, when compared to the unstimulated (DMSO) control with a non-parametric, paired Wilcoxon test). These responses were higher than those observed when S proteins were considered (p= 0.0040 and 0.0002 for childhood exposure and Shingles cohorts, respectively) (**Fig 2A).**

**FIG 2.**
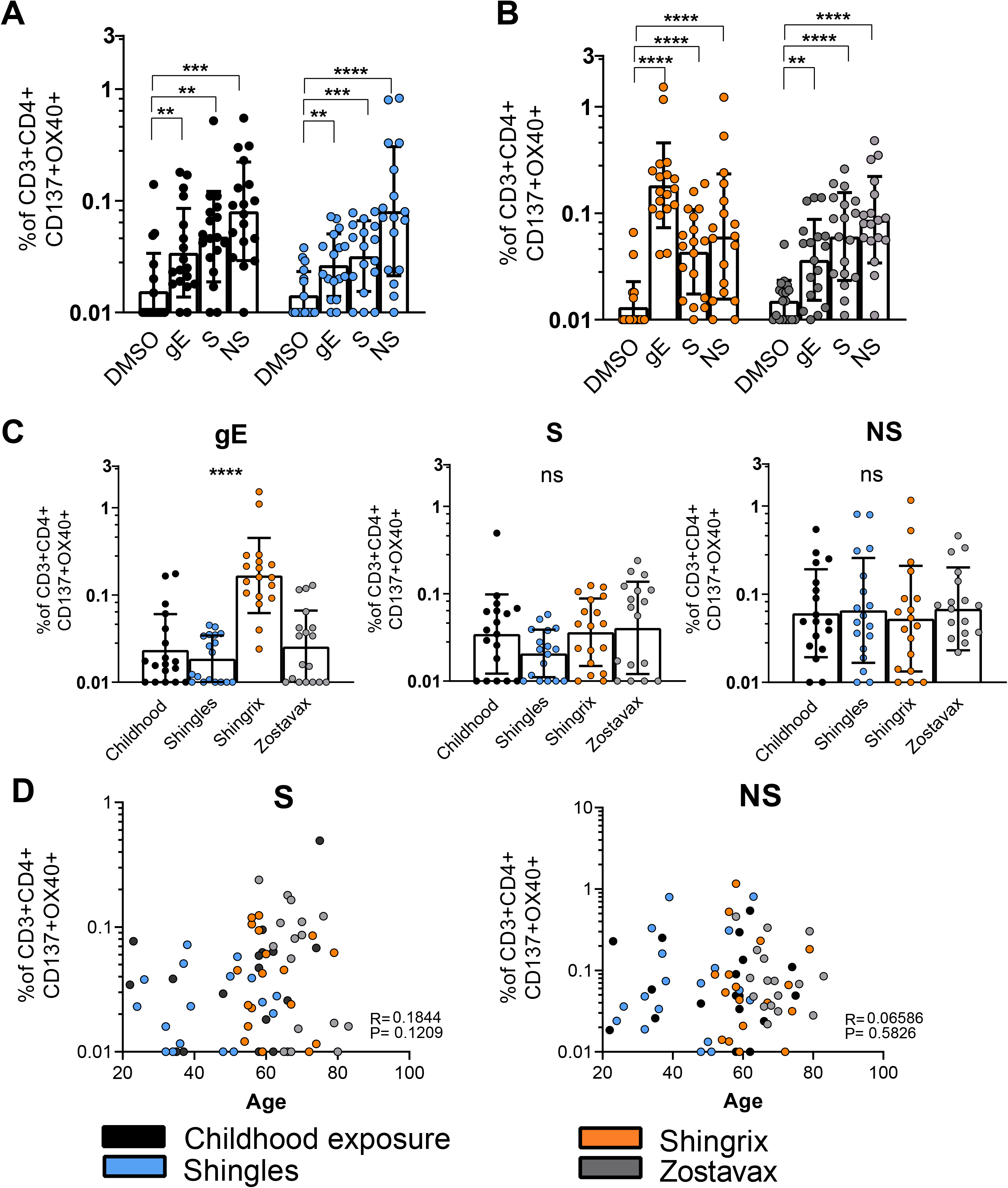
VZV-specific CD4+ T cell reactivity in childhood exposure, Shingles and HZ vaccinations. Frequency of CD137+OX40+CD4+ T cells (see **FIG 1** for gating strategy) was assessed for donor/MP stimulation combinations and compared to a paired DMSO negative control. A) Childhood exposure (black circles, n=18) and Shingles cohorts (blue circles, n=18). B) Shingrix (orange circles, n=18) and Zostavax (grey circles, n=18) cohorts. Data are expressed as the geometric mean and geometric standard deviation (SD). All reported statistical analyses have been performed using non-parametric, paired Wilcoxon test. C) Frequency of CD137+OX40+CD4+ T cells background subtracted and analyzed to compare the gE, structural (S) and non-structural (NS) MP across the cohort. Comparison across cohorts has been performed using Kruskal-Wallis test. D) Spearman correlation of the frequency of CD137+OX40+CD4+ T cells background subtracted based on donor Age after stimulation with structural (S) and non-structural (NS) MPs.

When the responses of the two vaccinated cohorts were examined, Zostavax and Shingrix vaccinees were found to be associated with significant responses to the various peptide pools. As expected, responses in Shingrix vaccinees were remarkably vigorous towards gE (Shingrix: GeoMean = 0.18, p <0.0001 compared to the unstimulated (DMSO) control) (**Fig 2B)**. These results confirmed that Shingrix vaccination induces a strong CD4+ T cell response directed against the gE antigen. We then investigated whether we could observe a difference in protein reactivity across the cohorts analyzed (**Fig 2C**). Comparisons across cohorts have been performed with background subtracted data and, interestingly, no systematically significant differences where observed for structural and NS protein pools (S: p=0.1491; NS: p=0.8708 by Kruskal-Wallis test) while, as expected, a significant difference across cohorts is observed in the case of the gE protein, driven by Shingrix vaccination (gE: p<0.0001 by Kruskal-Wallis test) (**Fig 2C**). When we specifically analyzed S and NS proteins in Shingrix and Zostavax vaccinees to determine whether these responses are towards similar epitope repertoires, no significant differences were observed (Shingrix Vs Zostavax S: p= 0.6337; NS: p= 0.4381, Mann Whitney test).

To understand if the similarity observed in protein reactivity across cohorts was due to the fact the cohorts had a significantly different age-range, we performed correlation of the CD4+T cell reactivity against structural and NS MPs with the age of the subjects, and found no significant correlation (Structural protein MP(S): r=0.1844 p=0.1209; non-structural proteins MP (NS): r=0.0659 p=0.5826) (**Figure 2D**).

### Immunodominant gE regions recognized specifically by Shingrix donors

Next, we defined the gE epitopes recognized in the Shingrix vaccinee cohort described above (**Table 1**), and performed donor HLA typing (**Table S2)** to enable determination of the HLA restriction of any responding epitope. As we wanted to ensure that our cohort was representative of the general population, we compared the frequencies of the main HLA class II alleles observed in our cohort with those observed in our repository of over 3,500 donors (**Fig 3A** and **Table 2**). This repository is inclusive of donors from a variety of clinical studies and representative of a diverse set of ethnicities and populations, ranging from the USA, South and Central America, Asia, South Africa and Europe, and therefore is a reasonable representation of the general population, with the further advantage of being HLA typed using the same methodology. Of the 28 different HLA class II alleles with phenotypic frequencies >10% in our cohort, 20 (71%) are also present in the general population with frequencies >10%. Additionally, 24 of the 27 (89%) alleles included in a reference panel of the most common and representative class II alleles in the general population (21) are also present in the current cohort (**Fig 3A**).

**FIG 3.**
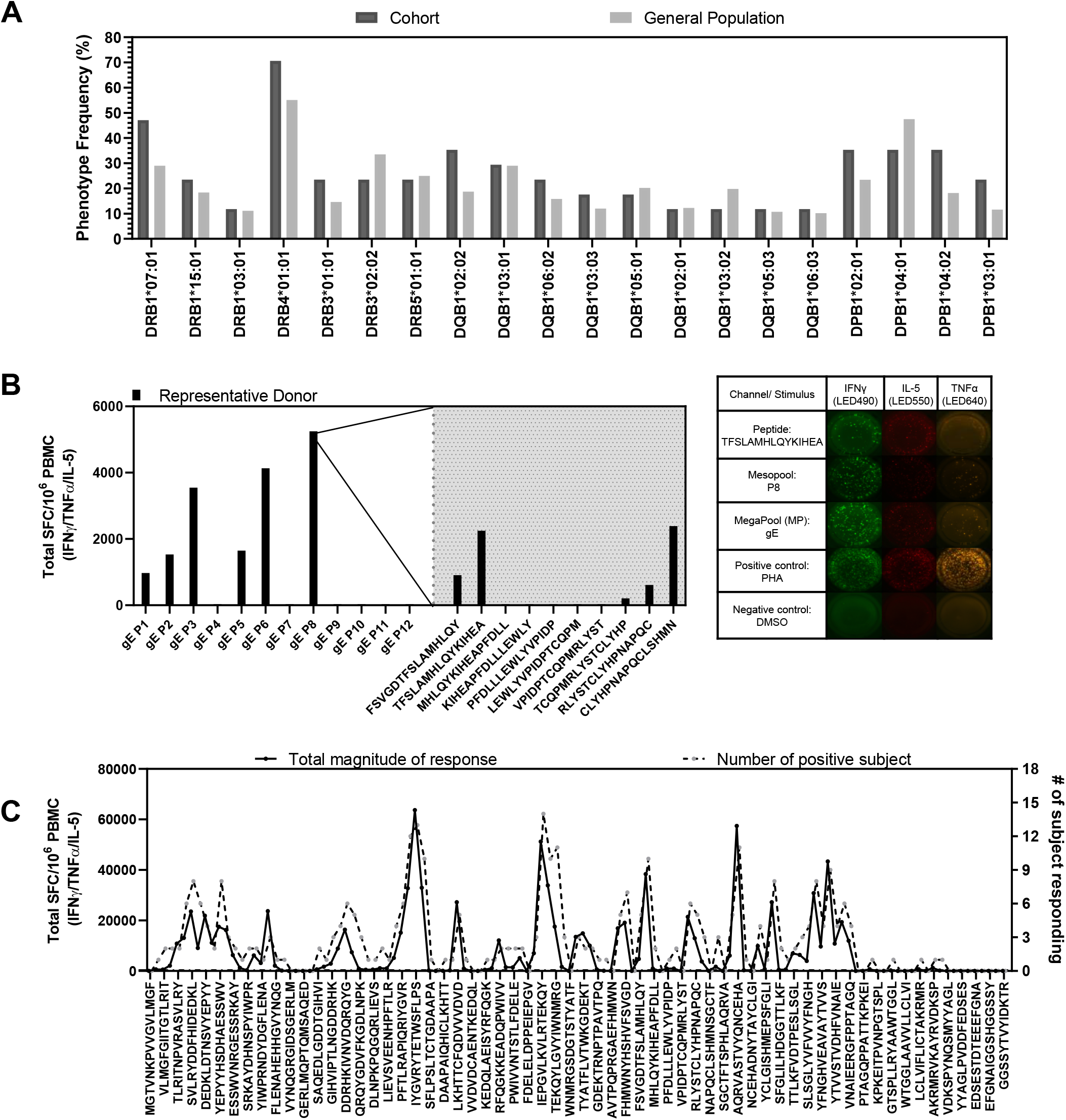
Epitope mapping of Shingrix vaccinees. A) HLA frequency of the Shingrix cohort (n= 18) compared to general population HLA frequency (n=3,500). B) Experimental scheme; representative donor with the mesopool reactivity at day 14 and peptide deconvolution of one of the positive mesopools at day 17 and relative FluoroSpot images for IFNγ, IL-5 and TNFα. C) Immunodominant regions of the gE protein showing the total magnitude of responses (solid lines) and the number of responding subjects (dotted lines).

**Table 2.**
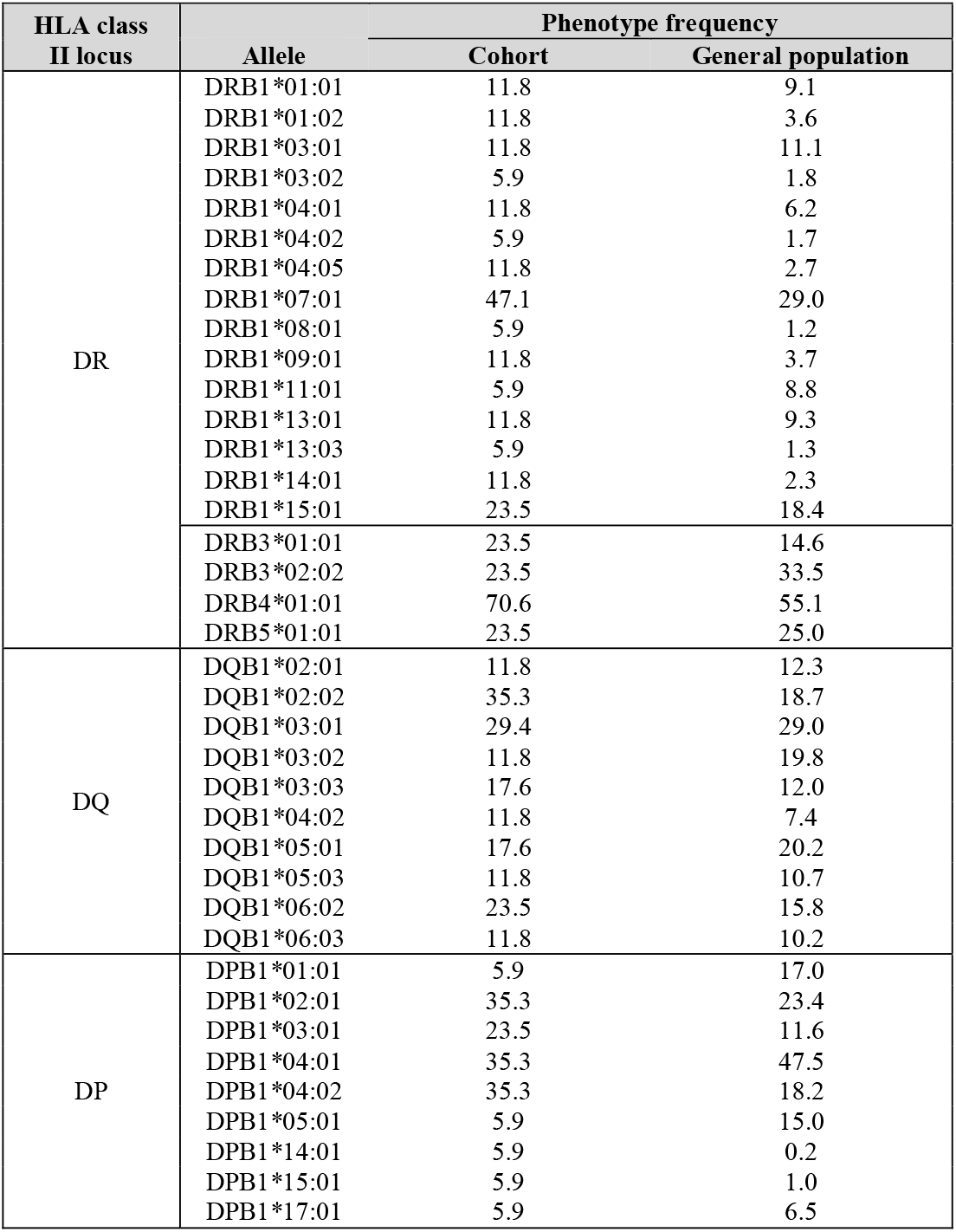
HLA class II allele frequency in Shingrix cohort compared to the general population.

The gE epitope identification strategy is summarized in Figure **3B-C**. To identify individual epitopes, PBMCs were expanded with the gE MP for 14-17 days. On day 14, the gE megapool and smaller pools composed by 10 or 11 peptide each (mesopools) were tested for cytokine reactivity. The individual peptides of the positively identified mesopools were tested for cytokine reactivity on day 17. On both days 14 and 17, reactivity was assessed using FluoroSpot assay (**Fig 3B**). It has been previously shown that the ASB01 adjuvant, in general, induces a mixed Th1/Th2 T cell phenotype (16). Therefore, to comprehensively identify the epitopes induced after Shingrix vaccination, while disregarding T cell polarization, we measured IFNγ, TNFα and IL-5 in the FluoroSpot assays by counting the single, double and triple producers only once, to avoid inflating our results.

Overall, we identified 89 epitopes recognized in at least one donor. When the response frequency and magnitude data were mapped to the gE sequence several immunodominant regions were identified (**Fig 3C**). Responses were found to be dispersed throughout the protein, with the exception of the 110 C-terminal residues, which were recognized poorly, if at all. A full list of the epitopes identified in this study has been submitted to the IEDB (http://www.iedb.org/1000857) and available in **Table S3**.

### Comparison with the repertoire of epitopes recognized in different donor cohorts

The data presented above in **Fig 2** indicate that Shingrix vaccinees have a stronger gE response as compared to other cohorts of VZV exposed/vaccinated individuals, and several epitopes were recognized per donor (**Fig 3C**). Those two points raised the question whether the higher Shingrix response was due to a stronger response to a similar number of epitopes, or responses of similar magnitude directed against a larger number of epitopes. To address this, we conducted pilot deconvolution experiments in two donors from the Zostavax cohort, and three donors from the childhood exposure cohort.

First, we inspected the overlap in repertoires between the Shingrix and Zostavax/childhood exposure donors. Of the 89 epitopes identified in the Shingrix cohort, 42 (47%) were also recognized in the Zostavax/childhood exposure samples. Conversely, only 5 (5%) of the peptides not recognized in Shingrix vaccinees were recognized in Zostavax/childhood exposure samples (p>0.0001 by the Chi-squared test) (**Fig 4A**). A Venn diagram is shown in **Fig 4B** to better illustrate instances of overlap and lack of overlap. This indicated that the epitope repertoire recognized in the Shingrix and Zostavax/childhood exposure donors is similar in nature, although Shingrix vaccination appears to induce a broader epitope repertoire for the gE protein. In support of this observation, Shingrix vaccinees responded to an average of 18±10 epitopes, with an average magnitude of 2041±1673 SFC/10^6^ PBMC per response. By contrast, the Zostavax/childhood exposure donors responded to 10 ± 8 epitopes, with a magnitude of 1353±1080 SFC/10^6^ PBMC per response. (**Fig 4C-D**). Thus, although these differences are not significant, Shingrix donors tend to have a broader epitope repertoire and responses of higher magnitude compared to the Zostavax/childhood exposure cohort.

**FIG 4.**
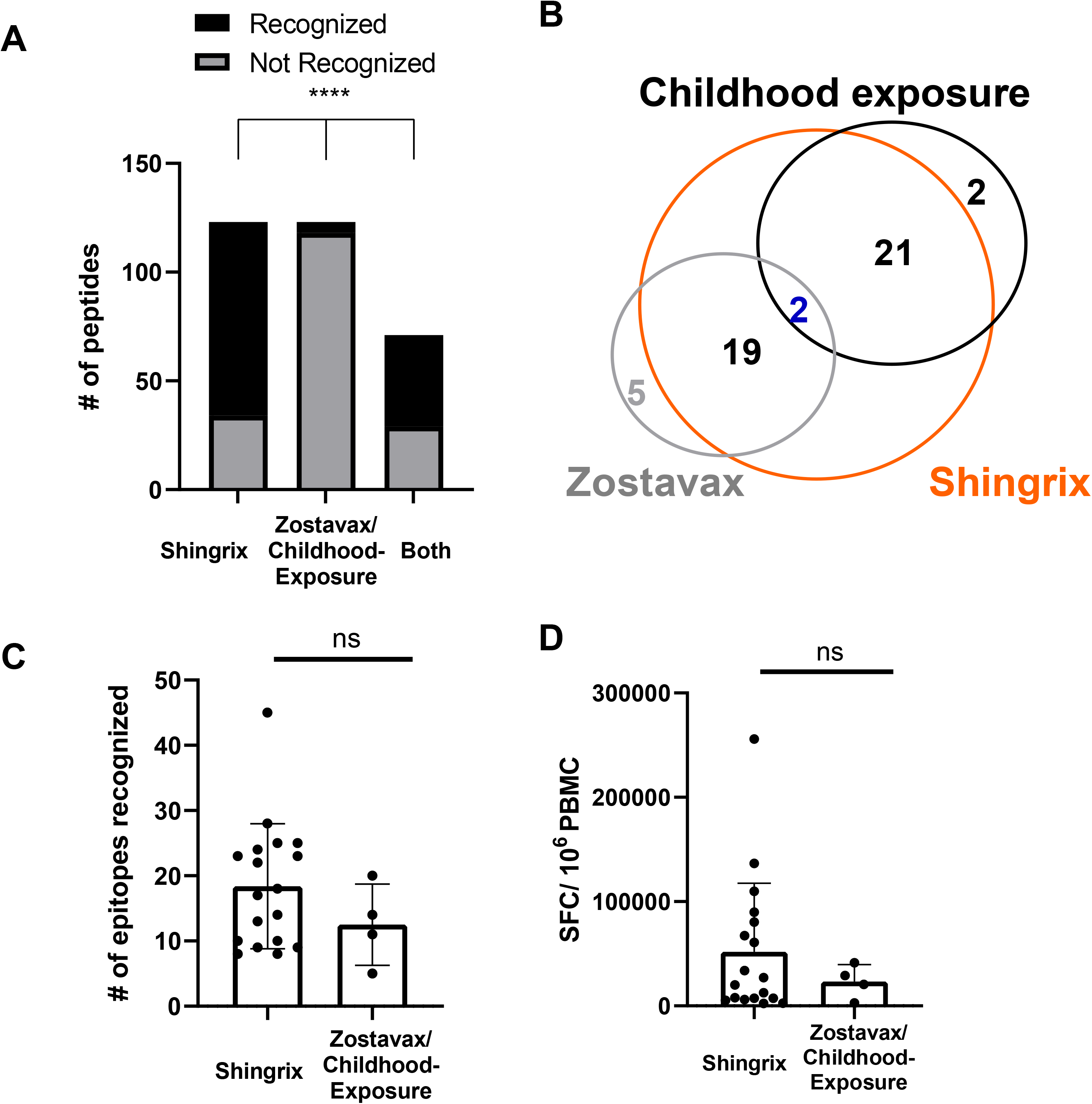
Shingrix patients show a trend for recognizing more epitopes and producing a stronger response. A) Number of recognized epitopes (black) over the non-recognized ones (grey) in Shingrix, Zostavax/childhood exposure or both (****P<0.0001, Chi-square test). B) Venn diagram showing epitope overlapping across Shingrix, Zostavax and Childhood exposure donors. C) Number of recognized epitopes per donor-cohort (ns; p= 0.3095) D) Magnitude of responses per donor-cohort (ns; p=0.7743). (C-D) Mann-Whitney test was applied for statistical comparisons across cohorts.

### Breadth and magnitude of gE-specific CD4+T cells

As mentioned above, of the 123 gE peptides tested, 89 were recognized in at least one Shingrix vaccinee, while only 34 were not recognized in any context (**Fig 4A**). Of the 89 epitopes inducing responses, 26 epitopes were recognized in only one donor, 19 epitopes were recognized in two donors, and the remaining 44 were recognized in three donors or more (**Fig 5B**). In terms of magnitude of response, the top 34 epitopes recognized in four donors or more were found to account for 80% of the total SFC response (**Fig 5C**).

**FIG 5.**
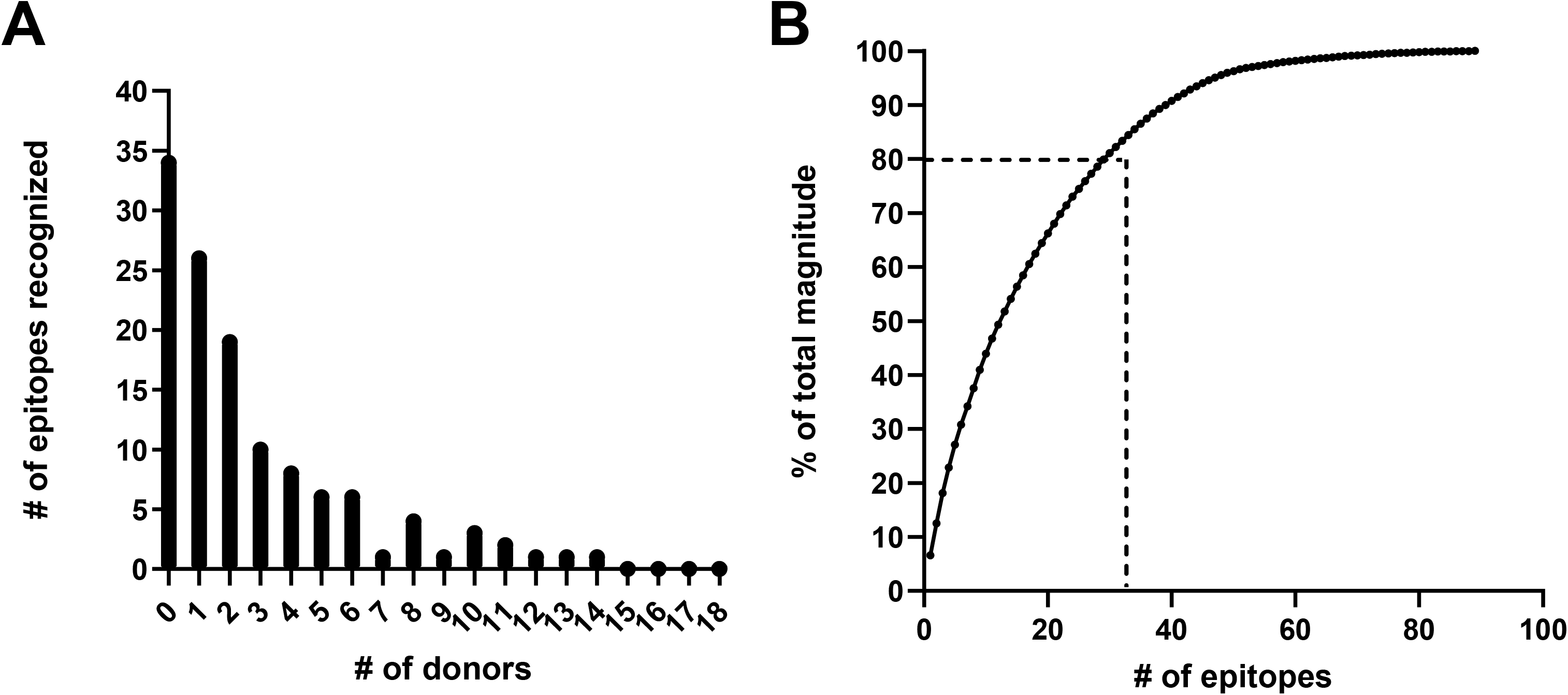
Shingrix vaccinees are associated with a broad CD4+ T cell response. A) The number of responding donors are plotted against the number of epitopes recognized. B) The total number of epitopes recognized are plotted against the percentage (%) of total magnitude of response (sum of total SFC of total responding donors).

**Table 3** lists these top 34 epitopes eliciting a response in four or more donors. The table also details the number of positive responses, the sum of peptide-specific SFC across donors, the corresponding percentage (%) of total SFC detected in the cohort, the cumulative SFC, and a response frequency score, calculated as described in the Methods. Overall, these data indicate that while Shingrix vaccinees are associated with a broad and diverse repertoire, a limited number of epitopes are dominant and account for the preponderance of the response. When we compared the 34 immunodominant epitopes characterized in Shingrix donors (**Table 3**), we found 7 were also recognized by Zostavax vaccinees, although they were not among the top most immunodominant epitopes.

**Table 3.**
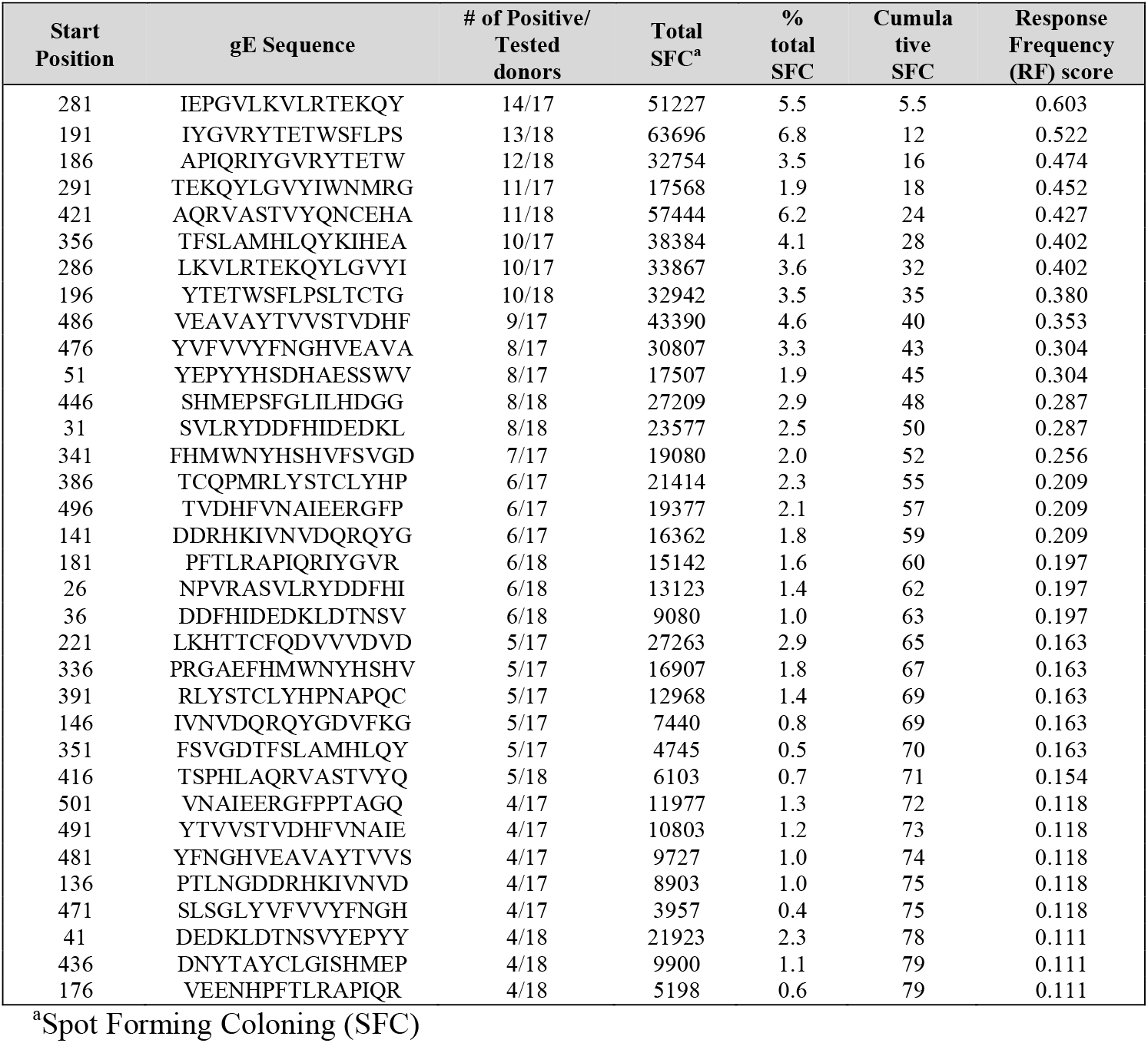
Most Commonly recognized gE epitopes among Shingrix vaccinees.

### HLA-binding capacity of dominant epitopes

To characterize the dominant epitopes in more detail we conducted HLA/epitope binding assays, using purified HLA molecules in vitro, as described in the Methods. **Table 4** presents an analysis of a set of 19 common HLA class II alleles selected on the basis of worldwide frequencies (21). Actual binding data was generated for 28 of 34 of the epitopes in **Table 3**. Six additional epitopes (136, 146, 176, 221, 471 and 481) were not available when the binding assays were performed, and in those cases, HLA binding predictions were used instead using the NetMHCIIpan algorithm (1). The use of HLA binding predictions for those peptides was supported by the strong correlation detected for predicted versus actual measured binding (**Fig 6**) in the cases where both experimental and predicted binding values were determined.

**Table 4.**
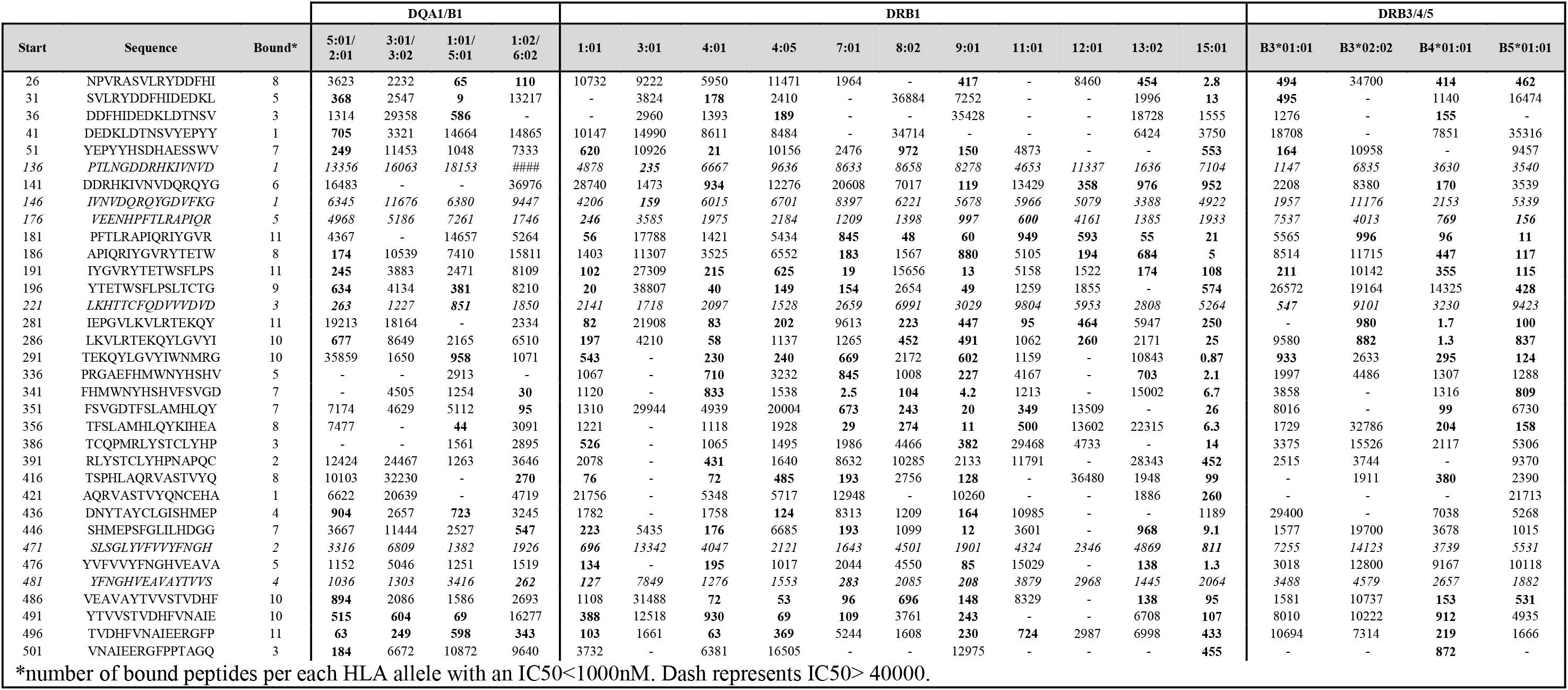
Observed in vitro binding capacity of 19 common HLA alleles to epitopes commonly recognized among Shingrix vaccinees. Predictive data is shown for italicized rows.*

**FIG 6.**
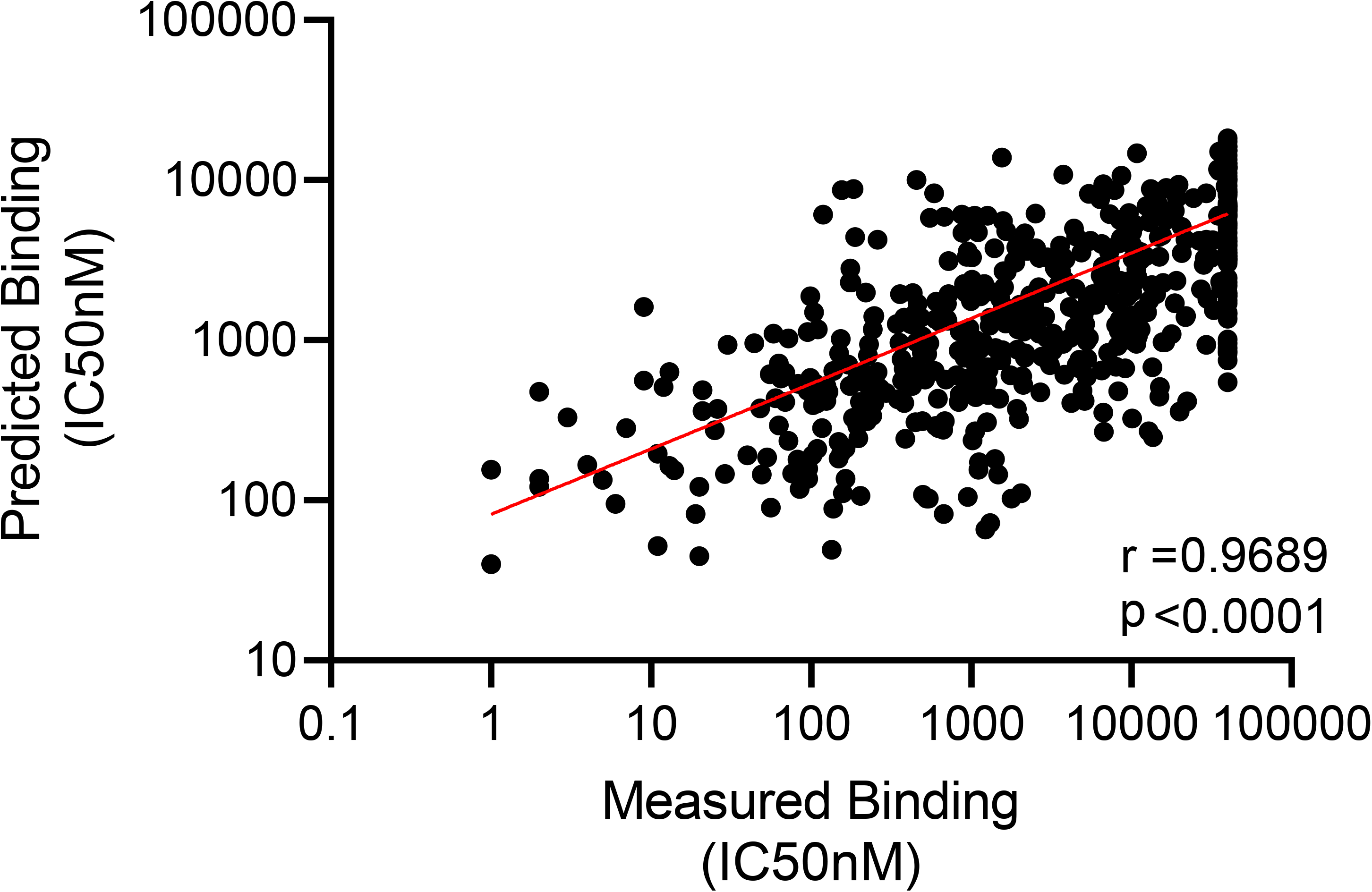
Correlation of predicted versus experimentally measured binding. Experimental HLA binding (x-axis, IC50 nM) compared to the predicted HLA binding (NetMHCpan 3.2, IC50nM,(1)). Pearson correlation was applied (r = 0.9689, p<0.0001).

Most of the dominant epitopes were found to be promiscuous HLA class II binders, with 50% binding to 7 or more alleles in the panel of 19 common specificities considered, and binding 6.1±3.4 alleles on average. By contrast, negative peptides (i.e. not recognized in any donor) bound an average of 2.1±3.6 alleles in the same panel of HLA variants.

### HLA restriction determinations

Next, we determined HLA restrictions for selected epitopes by the use of single HLA transfected cell lines. Briefly, T Cell Lines (TCL) generated by 14 day in vitro re-stimulation with an individual peptide were tested in antigen presentation assays, where different cell lines expressing a single HLA class II molecule were used as APCs (40), as detailed in the Methods. Representative results are shown **Fig 7A-C.** When a TCL derived from a donor responding to the gE 446-460 peptide (sequence SHMEPSFGLILHDGG) was tested with single HLA transfected APC lines matching the HLA type of the donor, a good response was observed in the case of the HLA DPB1*0402 APCs. No response was noted for any of the other APC lines matching the donor’s HLA type. Accordingly, the gE 446-460 peptide was determined to be DPB1*0402 restricted in this donor. Similarly, the gE 86-100 and 486-500 peptides (sequences NDYDGFLENAHEHHG and VEAVAYTVVSTVDHF, respectively) were identified as putatively restricted by DQB1*0302 and DRB1*0701 in other representative donors (denominated as “positive restrictions”). Conversely, the HLA/epitope combinations tested in the same assays and found negative were denominated as “negative restrictions”.

**FIG 7.**
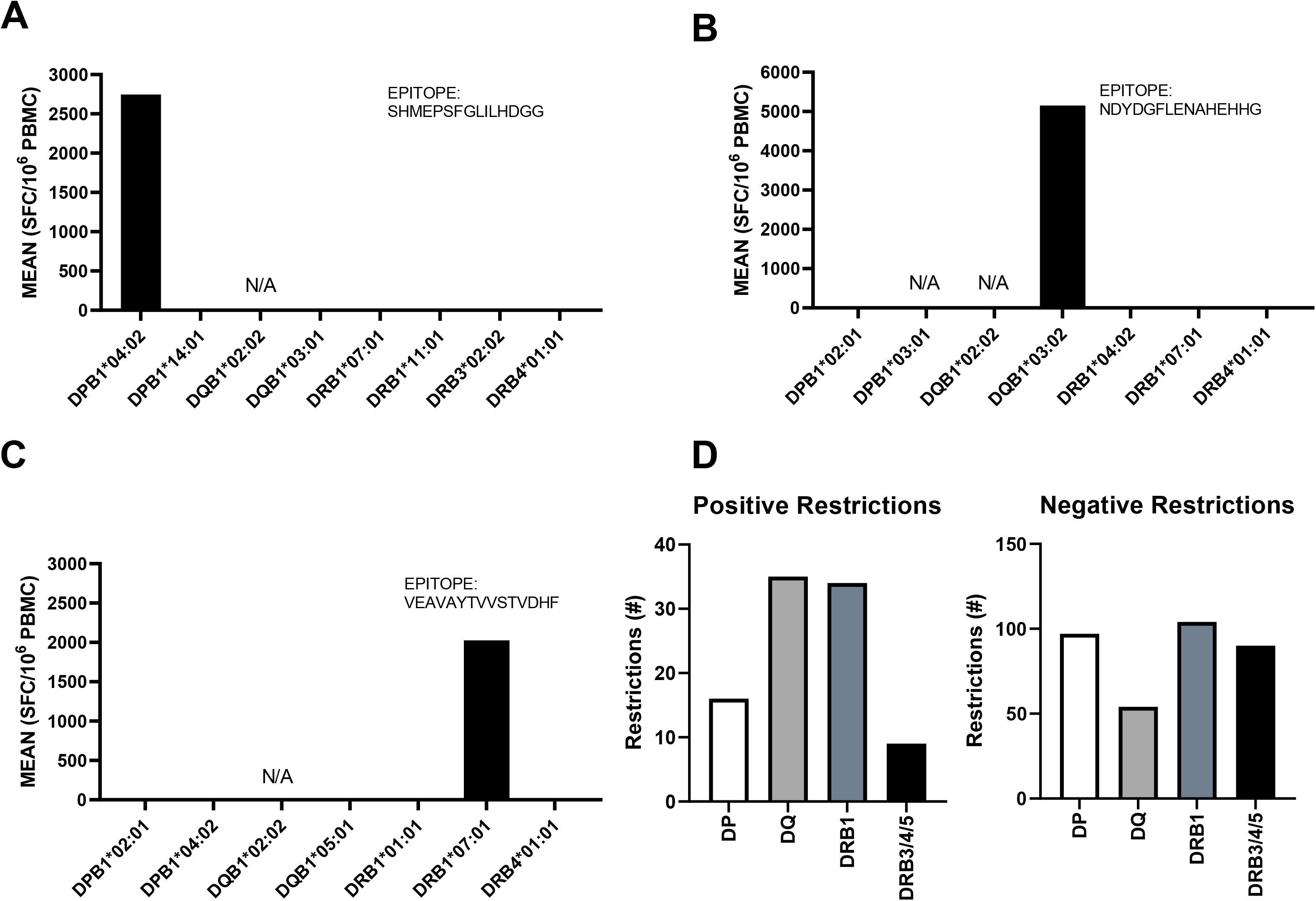
Donor HLA/epitope restrictions identified by HLA restriction assay. (A-C) Representative examples showing determination of HLA restriction for three epitopes in three donors. D) Summary of positive and negative epitope/HLA restrictions identified.

The results of the experiments are presented in **Table 5** and summarized in **Fig 7D**. Overall, we determined positive restriction for 38 peptides and 94 donor/peptide combinations (listed in **Table S3**). Most positive restrictions were associated with the DRB1 and DQ loci, consistent with previous studies (45). Not all restrictions could be determined unequivocally, as APC corresponding to some of the HLA alleles were not available because of limitations in cell availability or other technical issues. In addition to testing as many of the peptides in **Table 3** as possible, we also tested certain peptides that were positive in only one or few donors, but yielded vigorous responses. **Table 5** presents a summary of determined restrictions, including sequence, percentage of the response, number of donors recognized and associated HLA restriction(s).

**Table 5.**
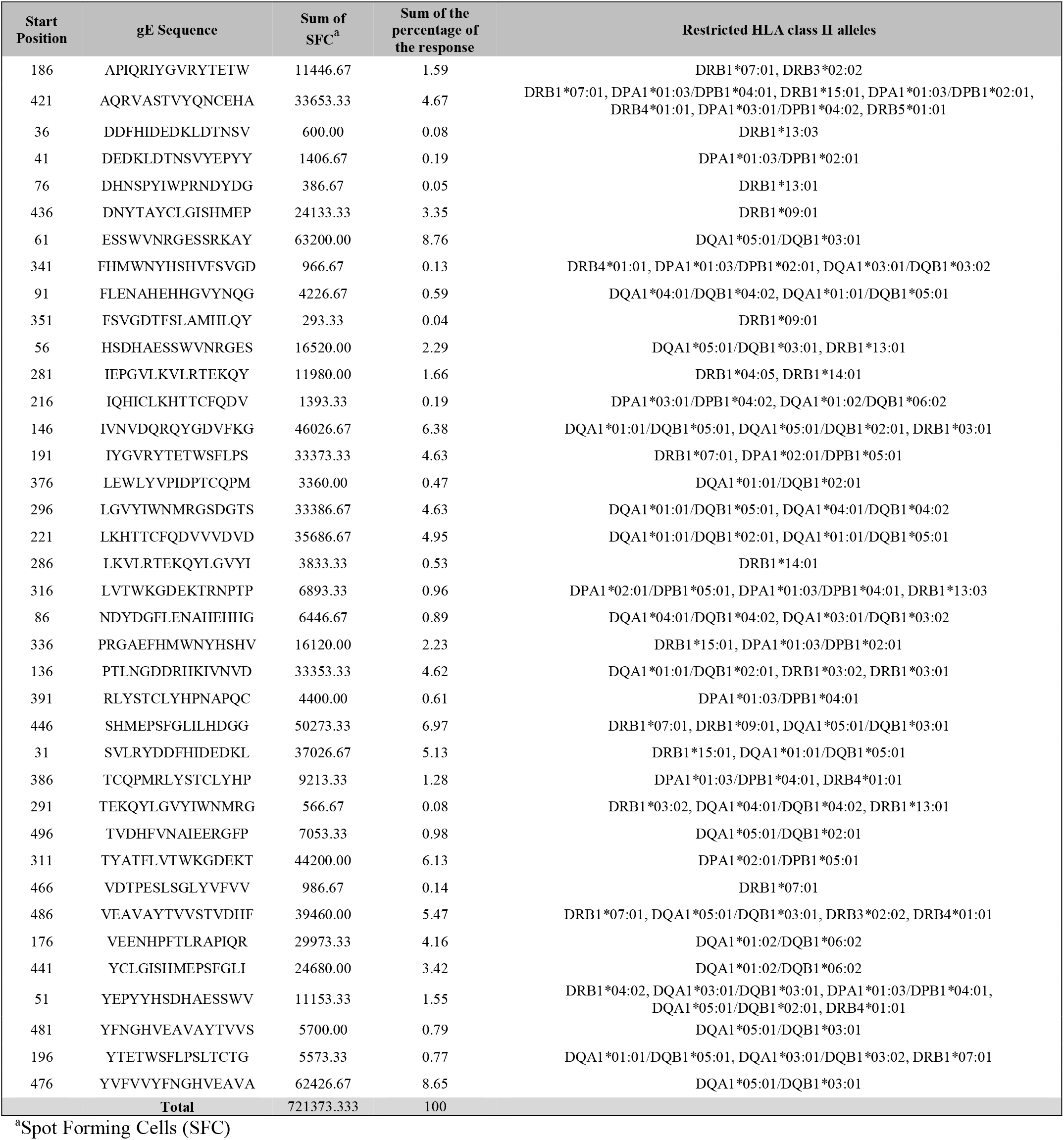
Summary of determined epitope/HLA restrictions.

### Predictive performance of different methodologies

The data set described above provided an opportunity to explore potential strategies to predict restrictions, without performing HLA restriction assays or performing in vitro HLA binding measurements. In a first determination, we explored the relative merit of four different methodologies to discern positive and negative restriction: the actual measured binding affinity, predicted binding affinity, percentile rank of predicted affinity, and the Response Frequency (RF) determination.

In terms of epitope prediction, different bioinformatics-based algorithms utilizing a broad array of computational approaches and platforms have been developed in the past 30 years. Several of the best performing algorithms, as determined by community benchmarking studies (51), are available in the IEDB Analysis Resource (IEDB-AR). Two different prediction metrics have been considered in this study: 1) predicted MHC binding affinity (or IC50 nM) and 2) relative percentile rank score, which is also based on the IC50 prediction, but ranks peptides as a function of their predicted affinity against a large random library of ligands, thereby allowing to identify the top percentage of epitopes, as required. While several studies have been performed to identify predicted affinity thresholds for individual MHC class I alleles, in general a prediction threshold of IC50 <500nM or percentile rank score of 2%, or better, have been found to cover >95% of the known class I epitopes; a thorough evaluation of MHC class II alleles and predictions is still ongoing (53).

To investigate this point further in the context of class II, we performed HLA predictions for the epitopes identified herein and the HLA class II alleles expressed in each Shingrix donor in our cohort, extracting both IC50 values and percentile scores, and used the results of the HLA restriction experiments to assess the efficacy of various prediction thresholds. An alternative approach that was considered was to identify predicted restriction by calculating the Relative Frequency of specific HLA alleles-epitope combinations identified across the Shingrix donors (48, 50). The Area Under the Curve (AUC) values observed in the receiver operating characteristic (ROC) curves were used to quantify predictive efficacy across the four methodologies analyzed.

As shown in **Fig 8A,** and as expected, actual measured binding was the most effective (AUC=0.7179), followed by prediction rank (AUC=0.6600). Since experimental binding determination is laborious and time consuming, for the subsequent analyses we focused on predicted rank as the most promising approach from the operational point of view. When the predictive efficacy was examined as a function of the different loci, we found that DRB1 (AUC= 0.7245; **Fig 8B)** and DQ (AUC=0.7542**; Fig 8C)** restrictions were predicted with reasonable accuracy. Conversely, the performance was much lower in the case of DP (AUC=0.5425; **Fig 8D**) and the DRB3/4/5 (AUC=0.6377; **Fig 8E**) loci, perhaps reflecting that these restrictions were relatively few and infrequent. Indeed, by considering only DRB1 and DQ loci as putative restriction elements, and utilizing the 15^th^ percentile rank as a threshold to define binding, we would have correctly called HLA restrictions for 39 of 94 instances, with 32 incorrect restrictions (false positives). In other words, utilizing those criteria we would be correct about 55% of the time and capture about 40% of the epitope restrictions.

**FIG 8.**
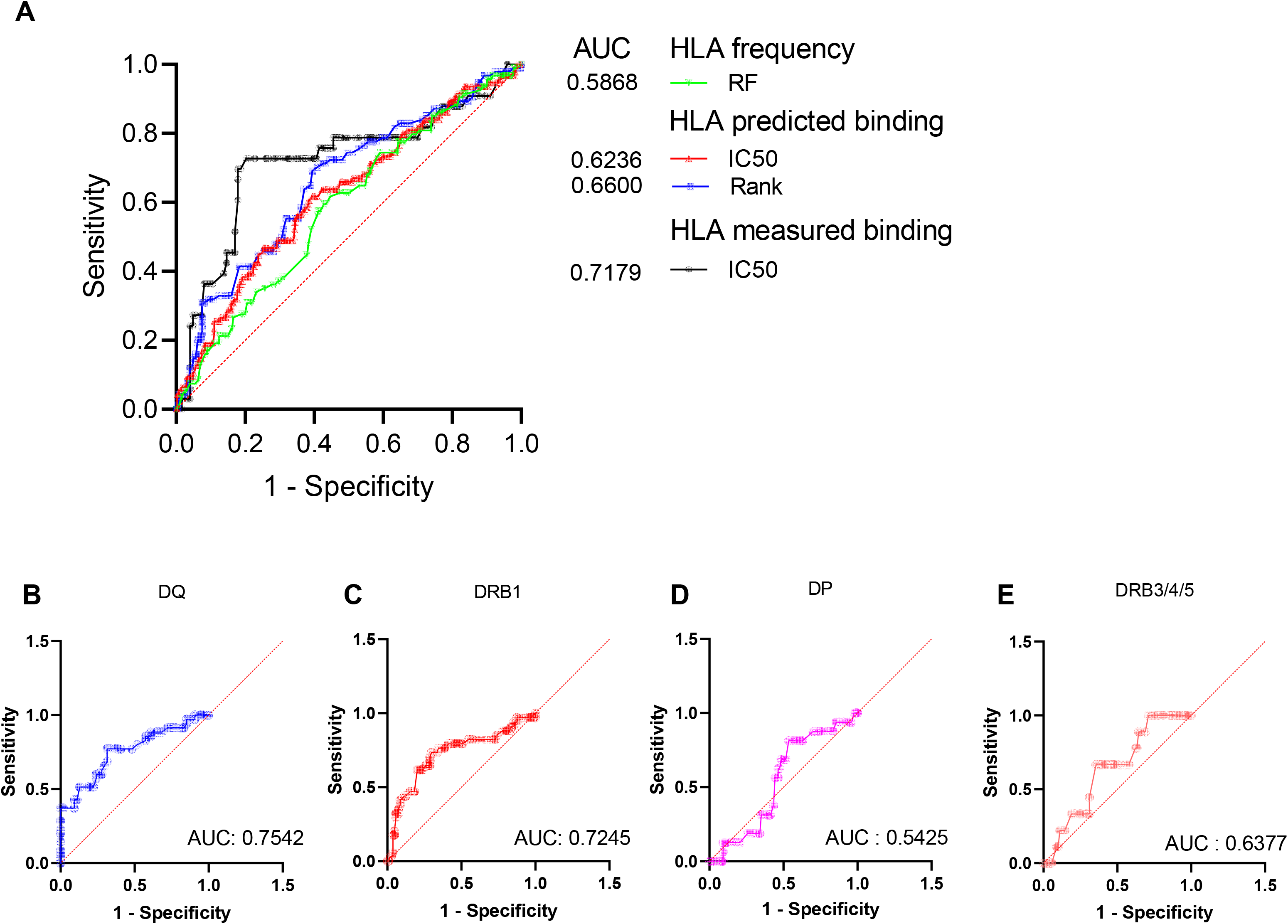
ROC curves of measured and predictive systems to infer HLA restriction. A) Comparison of different methodologies. B) HLA prediction based on rank percentile scores and as a function of the different HLA class II loci analyzed.

## Discussion

Here we report a detailed characterization of the epitope targets of the CD4 response of human subjects vaccinated with Shingrix. The results will enable mechanistic studies of vaccine efficacy, provide an example of a systematic analysis of epitope responses to a vaccine antigen, and have some methodological implications for future analyses and HLA restriction determinations.

Shingrix has shown increased efficacy in elderly and other groups as compared to the Zostavax vaccine (27, 54, 66). The reasons for this remarkable difference may involve intrinsic differences in the quality and durability of the responses elicited, also perhaps because of inclusion of the AS01B adjuvant. We believe that our results will facilitate mechanistic studies to directly address this point, by enabling HLA-class II tetramer studies and epitope pool-based isolation of reactive T cells.

In terms of enabling tetramer studies (11, 42), our results experimentally defined a total of 94 HLA/epitope restrictions, which allow to cover an overwhelming majority of the human population, irrespective of ethnicity. This set thus allows to overcome a limitation of studies that use tetrameric reagents to capture and study antigen specific T cells, namely that the tetrameric reagents generally available only cover limited allelic diversity, and are typically inclusive only of those alleles frequent in Caucasian populations. Our results also enable an alternative approach based on the use of pools of experimentally defined epitopes as “bait” to capture and characterize antigen specific T cells (38). Tetrameric reagents incorporating these epitopes may inform if and how specific epitopes affect polyfunctionality. Several studies have started to address these issues. Liang et al (32) reported that CD4+ T cells respond to peptides derived from different VZV ORFs, and that the response increases after Zostavax vaccination. Zostavax-specific CD4+ T cells have a mixed Th1/Th2/Th21 phenotype (36). Increases in T follicular helper cells (Tfh) and IL-21 in CD4s, in general, and a mixed Th1/Th2 response with both IFNγ and IL-4 release were reported from PBMCs (36). Levin et al. (34) reported that the profiles and magnitude of Th1 T cell memory responses differentiate Shingrix and Zostavax.

Shingrix vaccination, unsurprisingly, induces a vigorous gE-specific response, in agreement with previous studies (16, 34, 55). Interestingly, we were also able to detect a significant response against non-structural (NS) proteins, as well as other structural proteins, in Shingrix vaccinees. These responses are most likely derived by previous natural exposure or chickenpox vaccination. When we compared the responses across Zostavax vaccinees, childhood exposed and HZ patients were all found to have a similar response profile, with the strongest responses towards Non-structural (NS) proteins. This suggests also that the pattern of VZV specific CD4+ T cell protein immunodominance after natural exposure is in general against NS proteins. In this sense, Zostavax vaccination mirrors the natural infection profile, while Shingrix skews the natural protein immunodominance toward gE, the only component of the vaccine. Indeed, the T cell reactivity against gE induced by Shingrix is so strong that it is significantly higher than the non-structural (NS) counterpart. This is also in contrast with the heterogenic pattern of CD4+ T cell reactivity observed in young adults after chickenpox vaccination, where no clear immunodominance against NS proteins is observed. Instead, a quite diverse pattern of reactivity is observed in different donors, with responses against both structural and NS proteins, suggesting that priming by natural infection, as opposed to chickenpox vaccination in the young, induces a skewed response against NS proteins only, and might contribute to the reduced response against Zostavax observed in the elderly (18, 59).

We positively identified a total of 89 epitopes recognized across 18 Shingrix donors. Of these, the 34 most responsive capture 80% of the total (SFC) response against all identified epitopes. Thus, while Shingrix vaccinees are associated with a broad and diverse repertoire, a limited number of epitopes are dominant and account for the majority of responses. These epitopes will help build an immunogenic profile of VZV, as currently only 25 VZV gE-derived T cell epitopes recognized in humans have been reported in the published literature, as captured by the IEDB. Of the top 34 epitopes identified in our study, 7 (20%) nest or overlap with previously identified epitopes, while the remaining 27 (80%) are novel.

The results in terms of epitope breadth and coverage can be interpreted in the context of other studies from our laboratories that have used similar methodology to define the epitope repertoire of different antigens by scanning with sets of overlapping peptides. These studies included mycobacterial (37), pollen (47), cockroach (46) and dust mite allergens (28), and pertussis (46) and tetanus (8). The pattern of 20-30 epitopes encompassing the vast majority of the responses is generally applicable, and in this respect the gE antigen does not present unusual characteristics that might explain the remarkable vaccine efficacy. A lack of reactivity is generally observed in the terminal end of the gE protein in the Shingrix vaccinees analyzed in this study. This can be explained by the fact that those residues are not included in the recombinant protein used for the Shingrix vaccination to facilitate protein preparation.

Our results indicate that Shingrix vaccination might result in a marginally broader repertoire, as compared to Zostavax and natural exposure, and a somewhat higher magnitude of responses, with differences in epitope immunodominance hierarchy. Indeed, broader responses have been invoked as a correlate of protection in the case of HCV and other viral infections (30, 56, 61). However, in the case of Shingrix versus Zostavax, differences in the epitope breadth were not statistically significant and any differences should also be interpreted with caution, since the Shingrix samples were generally obtained closer to vaccination as compared to the Zoster cohort.

It is of interest to address whether the epitopes identified for gE are specifically targeted following Shingrix vaccination. The current dataset is only based on limited donors, and further studies will have to investigate this issue further. Also, the cohorts studied are reflective of the general volunteer donor pool that answered our recruitment in the San Diego area. Despite every effort made to recruit women and minorities, still there is a bias towards males and Caucasians. As a further caveat, it should be considered that the cohort size utilized in the current study is somewhat limited. For all of the above reasons, the results should be interpreted with caution, and larger studies will be required to confirm our conclusions.

Finally, our data, in line with previous studies (38, 40), validate the use of single HLA transfected target cells as a valid methodology to experimentally determine HLA restrictions. The methodology is powerful and accurate, but somewhat laborious. For this reason, we explored the possibility of applying publicly available prediction approaches to infer the HLA-restriction of experimentally defined epitopes. In this context, our data set also allowed evaluation of the efficacy of three different methodologies to predict HLA restrictions in absence of experimental data. We found that the percentile rank metric is the best approach to predict restrictions for the DRB1 and DQ loci. Using a 15^th^ percentile cut-off, we were able to accurately predict 40% of epitope/HLA combinations for DRB1 and DQ alleles identified in our dataset. This conclusion has two potential applications: the first is to assign, with high likelihood, the HLA-restriction of experimentally defined epitopes, and the second is to apply prediction algorithms before running experiments to selectively test predicted candidates only in the specific donor(s) with corresponding HLA type. This second application can help narrow down targets of investigation prior to performing assays and increase the speed and coverage of identifying HLA/epitope combinations, while also conserving precious biological samples.

## Materials and Methods

### Study subjects, PBMCs isolation, and HLA typing

The Shingrix vaccine cohort consists of healthy donors; adult male and non-pregnant female volunteers, age 50 and over, that were enrolled after receiving Shingrix vaccination (n=18) under the LJI program VD-184. PBMC derived from Shingrix cohorts were processed at LJI by density-gradient sedimentation using Ficoll-Paque (Lymphoprep; Nycomed Pharma, Oslo, Norway). Isolated PBMC were cryopreserved in heat-inactivated fetal bovine serum (FBS; HyClone Laboratories, Logan UT), containing 10% dimethyl sulfoxide (DMSO) (Gibco) and stored in liquid nitrogen until use in the assays.

HLA typing was performed by an ASHI-accredited laboratory at Murdoch University (Western Australia) for Class II (DRB1, DRB3/4/5, DQA1/DQB1, DPB1) as previously described (22, 24, 38). **Table 1** describes general cohort characteristics including ethnicity, race, gender, age. **Table S2** lists the HLA-typing of the Shingrix vaccinees.

### Peptides, Mesopools, and Megapool

Overlapping 15-mers by 10 were derived from gE protein of the Dumas reference sequence (NC_001348.1) available in the Viral Expasy Database (https://viralzone.expasy.org/), for a total of 123 peptides. Peptides were synthesized as crude material (A&A, San Diego, CA) and resuspended in DMSO at a concentration of 20 mg/mL.

Part of the peptide synthesis was either pooled in small mesopools, containing 10 peptides each, or pooled together with all the gE peptides and sequentially lyophilized as previously described to generate a gE Megapool (3). gE epitopes identified in this study were mapped to the gE Dumas reference sequence (NC_001348.1) to identify immunodominant regions, using the Immunobrowser tool (15).

### Activation Induced Marker (AIM) Assay

Cells were cultured in the presence of either gE, structural (S), non-structural (NS) MPs (2ug/mL), DMSO (2%), or phytohemagglutinin (10ug/mL) (PHA) for 22-24 hours. After stimulation, cells were stained with surface markers for 30 minutes at 4°C. Detailed information for all the antibodies used for flow cytometry experiments in this study can be found in **Table 6.** Surface marker proteins were quantified via flow cytometry (LSRII, BD Biosciences) and analyzed using FlowJo software version 10.5.3 (TreeStar Inc., Ashland, OR). The gating strategy is schematically represented in **Fig 1**. Statistical analyses were performed using Graphpad Prism v.8 (San Diego, CA). Responses of paired samples across different stimuli were tested using a Wilcoxon matched paired test.

**Table 6.** AIM Flow Cytometry Panel. Antibody panel used in flow cytometry experiments to identify both magnitude and quality of CD4+ T Cell response.

| Antibody | Flouorochrome | Volume (µl/well) | Vendor | Catalog | Clone |
| --- | --- | --- | --- | --- | --- |
| CD3 | AF700 | 2 | Biolegend | 317340 | OKT3 |
| CD14 | V500 | 1 | BD | 561391 | M5E2 |
| CD19 | V500 | 1 | BD | 561121 | HIB19 |
| Live/Dead | ef506/Aqua | 0.5 | eBioscience | 65-0866-18 | -- |
| CD8 | Cy5.5 | 2 | eBioscience | 45-0088-42 | RPA-T8 |
| CD4 | Pacific Blue | 2 | BD | 558116 | RPA-T4 |
| OX40 | PE-Cy7 | 2 | Biolegend | 350012 | Ber-ACT35 |
| CD137 | APC | 2 | Biolegend | 309810 | 4B4-1 |

### Epitope identification by FluoroSpot

PBMCs were expanded in the presence of gE Megapool (MP) at a concentration of 1μg/mL for 14 days. After culturing, cells were collected and restimulated in triplicate at a density of 5×10^4^ cells/well with gE MP (1μg/mL), smaller mesopools containing 10 peptides each (1μg/mL), PHA (10μg/mL), and DMSO (0.1%) in 96-well plates previously coated with anti-cytokine antibodies for IFNγ, TNFα, and IL-5 (mAbs 1-D1K, MT25C5, TRFK5, respectively; Mabtech, Stockholm, Sweden) at a concentration of 10μg/mL.

After 20hr incubation at 37°C, 5% CO2, cells were discarded and FluoroSpot plates were washed and further incubated for 2hr with cytokine antibodies (mAbs 7-B6-1-BAM, 5A10 biotinylated, MT20D9-WASP, respectively; Mabtech, Stockholm, Sweden). Subsequently, plates were washed again with PBS/0.05% Tween20 and incubated for 1hr with fluorophore-conjugated antibodies (Anti-BAM-490, SA-550, and Anti-WASP-640). Computer-assisted image analysis was performed by counting fluorescent spots using an AID iSPOT ELISPOT reader (AIS-diagnostika, Germany).

Each mesopool was considered positive compared to the background based on the following criteria: 100 or more spot forming cells (SFC) per 10^6^PBMC after background subtraction for each cytokine analyzed, a stimulation index (S.I.) greater than 2, and statistically different from the background (p<0.05) in either a Poisson or T test. Mesopools eliciting the strongest cytokine response were deconvoluted at day 17 into the corresponding peptides at a concentration of 10ug/mL, and analyzed using the same criteria applied for the mesopools.

### Single HLA transfected cell lines

Single HLA transfected fibroblast (DAP.3) or B lymphocyte (RM3) cell lines were generated and maintained as previously described (38, 40). Based on the HLA typing of the donor, we assigned the HLA transfected cells considering only matches for the HLA class II DR, DP and DQ beta chains. When multiple cell lines correspond to the same beta chain, the alpha chain was matched based on the typing as well.

Short term T Cell cultures (TCL) were set up using donor PBMC cultured individually in the presence of single epitopes previously identified in the epitope identification experiments at 1ug/mL concentration for 14 days. At day 13, APC lines fibroblast-derived were induced with butyric acid (100 μg/mL) to favor HLA-expression on the cell membrane.

The following day, all APC lines were harvested, and viability (all >75%) was determined using Trypan Blue. Each APC line was pulsed individually with peptide at a concentration of 10ug/mL for 1hr at 37 °C, 5% CO_2_. TCLs were stimulated in triplicate with either a peptide-pulsed APC line, APC line alone (as a control), peptides (1μg/mL), PHA (10μg/mL), or DMSO (0.1%) in 96 well plates previously coated with anti-cytokine antibodies for IFNγ, TNFα, and IL-5 at a concentration of 10μg/mL. FluoroSpot assays and analysis criteria were carried out as described above for the epitope identification experiments. Computer-assisted image analysis was performed by counting fluorescent spots using Mabtech’s IRIS FluoroSpot and ELISpot reader (Mabtech, Sweden).

### HLA binding assay

Classical competition assays to quantitatively measure peptide binding to purified class II MHC molecules were performed as previously described (58). Briefly, a high affinity radiolabeled peptide was co-incubated at room temperature or 37C with purified MHC, a cocktail of protease inhibitors, and various concentrations of unlabeled inhibitor peptide. Following a two-day incubation, MHC bound radioactivity was determined by capturing MHC/peptide complexes on Ab coated Lumitrac 600 plates (Greiner Bio-one, Frickenhausen, Germany), and measuring bound cpm using the TopCount (Packard Instrument Co., Meriden, CT) microscintillation counter. The concentration of peptide yielding 50% inhibition of the binding of the radiolabeled peptide was calculated. Under the conditions utilized, where [label]<[MHC] and IC50 ≥ [MHC], the measured IC50 values are reasonable approximations of the true Kd values. Each competitor peptide was tested at six different concentrations covering a 100,000-fold range, and in three or more independent experiments. As a positive control, the unlabeled version of the radiolabeled probe was also tested in each experiment.

### RF score calculation and HLA epitope prediction

T cell epitope prediction analyses, as well as Relative Frequency (RF) score calculations, were performed using IEDB analysis tools (14). The relative frequency (RF) score is a combined metric that accounts for the positivity rate (how frequently an epitope elicits a positive response) and the number of independent assays. Specifically, RF = [(r − sqrt(r)]/t, where r is the total number of responding donors and t is the total number of donors tested (3). The RF score has been calculated using RATE tool (48, 49). HLA class II binding predictions were performed following two different strategies: one using NetMHCIIpan 3.2 (29) only and extracting IC50 predicted values, and the other one using the IEDB recommended 2.22 methodology and extracting rank percentile values. The IEDB recommended 2.22 methodology is based on a combination of seven prediction algorithms with priority on the consensus method (65), which includes comblib (57), SMM (44) and NN (43) algorithms, followed by NetMHCIIpan 3.2 (29).

## Data Availability

A complete list of the epitopes identified in this study has been submitted to the IEDB (http://www.iedb.org/1000857).

## Acknowledgments

We would like to thank the LJI flow cytometry core facility for their outstanding expertise. We thank Jose Barrera, Sydney Dong, Brittany Schwan, and Gina Levi for coordinating sample collection and processing. We thank Connor Kidd for his assistance carrying out experiments. We thank Bali Pulendran, Rafi Ahmed, Shane Crotty and Nadine Raphauel for helpful discussions and suggestions. Research reported in this publication was supported by the National Institute of Allergy And Infectious Diseases of the National Institutes of Health under Award Number U19AI142742. The content is solely the responsibility of the authors and does not necessarily represent the official views of the National Institutes of Health. All the authors declare no conflict of interest.

H. Voic, R. de Vries, P. Rubiro, and E. Moore performed experiments. R. de Vries and J. Sidney assisted in the bioinformatics analyses. S. Mallal and E. Philips performed HLA typing. B. Schwann provided samples and sample information. A. Grifoni and H. Voic reviewed the data and planned the experimental strategy. H. Voic, A. Grifoni, A. Sette and D. Weiskopf conceived and directed the study and wrote the manuscript. All the authors have critically read and edited the manuscript.

## Notes

### Competing Interest Statement

The authors have declared no competing interest.

### Summary of Updates

-Fix issues due to incorrect merging of multiple coauthors versions. -Updated Figures order -Updated Figure 2 -Updated results description and data discussion

